# Methylation Data Processing Protocol & Comparison of Blood and Cerebral Spinal Fluid Following Aneurysmal Subarachnoid Hemorrhage

**DOI:** 10.1101/2020.03.24.005264

**Authors:** Annie I. Arockiaraj, Dongjing Liu, John R. Shaffer, Theresa A. Koleck, Elizabeth A. Crago, Daniel E. Weeks, Yvette P. Conley

## Abstract

One challenge in conducting DNA methylation-based epigenome-wide association studies (EWAS) is the appropriate cleaning and quality-checking of the methylation values to minimize biases and experimental artifacts, while simultaneously retaining potential biological signals. These issues are compounded in studies that include multiple tissue types, and/or tissues for which reference data are unavailable to assist in adjusting for cell-type mixture, for example cerebral spinal fluid (CSF). For our study that evaluated blood and CSF taken from aneurysmal subarachnoid hemorrhage (aSAH) patients, we developed a protocol to clean and quality-check genome-wide methylation levels and compared the methylomic profiles of the two tissues to determine whether blood is a suitable surrogate for CSF. CSF samples were collected from 279 aSAH patients longitudinally during the first 14 days of hospitalization, and a subset of 88 of these patients also provided blood samples within the first two days. Quality control (QC) procedures included identification and exclusion of poor performing samples and low-quality probes, functional normalization, and correction for cell-type heterogeneity via surrogate variable analysis (SVA). Significant differences in rates of poor sample performance was observed between blood (1.1% failing QC) and CSF (9.12% failing QC; p = 0.003). Functional normalization increased the concordance of methylation values among technical replicates in both CSF and blood. Likewise, SVA improved the asymptotic behavior of the test of association in a simulated EWAS under the null hypothesis. To determine the suitability of blood as a surrogate for CSF, we calculated the correlation of adjusted methylation values between blood and CSF globally and by genomic regions. Overall, mean correlation (r < 0.26) was low, suggesting that blood is not a suitable surrogate for global methylation in CSF. However, differences in the magnitude of the correlation were observed by genomic region (CpG island, shore, shelf, open sea; p < 0.001 for all) and orientation with respect to nearby genes (3’ UTR, transcription start site, exon, body, 5’ UTR; p < 0.01 for all). In conclusion, the correlation analysis and QC pipelines indicated that DNA extracted from blood was not, overall, a suitable surrogate for DNA extracted from CSF in aSAH methylomic studies.

## 1 Introduction

The epigenome-wide association study (EWAS) approach has emerged in recent years as a hypothesis-free method for investigating the associations between epigenetic marks, such as DNA methylation, and human phenotypes. Challenges pertaining to the cleaning and processing of methylomic data persist, including issues related to sample quality, controlling for cell type heterogeneity, comparing methylomic profiles across tissue types, and modeling dynamic changes in methylation over time. Here, we describe our quality control (QC) pipeline for processing and quality-checking genome-wide methylation data obtained from samples of blood and cerebral spinal fluid (CSF) in a cohort of acute subarachnoid hemorrhage (aSAH) patients. aSAH is a form of stroke leading to variation in clinical outcomes such as cerebral vasospasm, coma, delayed cerebral ischemia (DCI), cognitive decline, and death (Wermer et al. 2007). Previous work (Endres et al. 2000; Nelson, Kavalali, and Monteggia 2008; Stapels et al. 2010) has suggested that changes in DNA methylation occur following aSAH. We hypothesize that these methylomic changes may be clinically relevant. Therefore, the overreaching goal of this ongoing initiative is to understand the changes in methylomic profiles occurring after aSAH to identify biomarkers predictive of prognosis and recovery outcomes. The purpose of this specific study was to develop and implement a pipeline for cleaning and quality-checking methylomic profiles derived from CSF tissue and to determine the suitability of peripheral blood as a surrogate for CSF.

## 2 Materials and Methods

### 2.1 Study Design Overview

Our study population is comprised of individuals who have sustained an aSAH. Patient DNA was obtained from two biological tissues, CSF (drained as standard of care) and blood. This study investigated CSF samples collected longitudinally from 279 patients during the first 14 days of hospitalization, and blood samples from 88 of these individuals collected within the first day of hospitalization. Methylomic profiles were obtained using a genome-wide array, from which methylation levels, quantified as beta-values (i.e., percent methylation) and M-values (i.e., a transformation of the beta-values, which exhibit beneficial properties for statistical analysis), were assessed for over 450,000 cytosine-phosphate-guanine (CpG) sites. QC analyses of methylation data were performed in the R statistical computing environment using the following packages: minfi (Aryee et al. 2014), ENmix (Xu et al. 2016) and sva (Leek et al. 2012). After QC, cleaned methylomic profiles were contrasted between blood and CSF samples to determine the utility of blood as surrogate for CSF.

### 2.2 Patient Recruitment and Sample Collection

Participants were considered for this study if they were admitted to the University of Pittsburgh Medical Center Neurovascular Intensive Care Unit with an aSAH confirmed by digital subtracted cerebral angiography and/or head computed tomography (CT) and a Fisher grade (measure of hemorrhage burden) > 1. Informed consent was obtained from the participant or their legal proxy using a protocol approved by the University of Pittsburgh Institutional Review Board. Exclusion criteria included a history of debilitating neurologic disease or subarachnoid hemorrhage due to arteriovenous malformation, trauma, or mycotic aneurysm.

Daily CSF samples were collected for the first 14 days after aSAH from an external ventricular drain placed as standard of care and DNA extracted using the Qiamp Midi kit (Qiagen, Valencia, CA, USA). Venous blood was collected within the first day of hospitalization and DNA was extracted using a simple salting out procedure. All DNA was stored in 1X TE buffer at 4°C.

This study included 279 aSAH patients. For the CSF samples, we targeted days 1, 4, 7, 10, and 13 post-aSAH, and substituted samples +/- 1 day when target days were unavailable. Blood samples collected within the first day of hospitalization after aSAH were included in this study for 88 of the 279 participants.

#### 2.2.1 Potential Covariate Assessments

The severity of aSAH was assessed by Fisher grade (Fisher, Kistler, and Davis 1980) employing CT scan to assess hemorrhage burden and by Hunt and Hess scores (Hunt and Hess 1968) to assess symptom burden. Demographic and anthropometric characteristics such as age, sex, race, height, and weight were collected from medical records (Table 1). Smoking status was also collected.

**Table 1.** Characteristics of study participants.

|  | Blood (N=88) | CSF (N=279) |
| --- | --- | --- |
| Demographic |  |  |
| Age (years) | 52.4±11.1 | 52.9±11.0 |
| Gender (female/male) | 59/29 (67%) | 193/86 (69%) |
| Race (white/black/others) | 76 (86%)/ | 243 (87%)/ |
|  | 10 (11%)/ | 30 (11%)/ |
|  | 2 (2%) | 6 (2%) |
| Height (in) | 66.1±4.8 | 66.1±4.3 |
| Weight (kg) | 80.6±19.7 | 78.6±20.0 |
| Clinical |  |  |
| Current smoke (yes/no) | 50/37 (57%) | 155/120 (56%) |
| Fisher grade (2/3/4) | 23 (26%)/ | 83 (30%)/ |
|  | 48 (55%)/ | 138 (49%)/ |
|  | 17 (19%) | 58 (21%) |
| Hunt & Hess score (1/2/3/4/5) | 2 (2%)/ | 20 (7%)/ |
|  | 26 (30%)/ | 84 (30%)/ |
|  | 37 (42%)/ | 109 (39%)/ |
|  | 19 (22%)/ | 48 (17%)/ |
|  | 4 (5%) | 18 (6%) |
| DCI (yes/no) | 45/42 (52%) | 123/154 (44%) |

Values are presented as mean±sd for continuous variables, category count followed by percentage of the first category for binary variables, and count (percentage) for variable with multiple categories. Missing values exist for some variables, accounting for the discrepancy between count sum and sample size.

### 2.3 DNA Methylation Data Collection and Plate Design

The Illumina (San Diego, CA) Infinium HumanMethylation450 BeadChip platform was used to assess the methylation levels at over 450,000 CpG sites in the samples. Methylation data collection was performed by the Center for Inherited Disease Research (CIDR) of Johns Hopkins University. Each BeadChip, hereafter referred to as a plate, consists of eight chips of 12 samples arranged in a layout of six rows by two columns. This enables 96 samples to be run on a single plate. To avoid plate effects, all blood samples were assayed together on a single plate. CSF samples were placed across 11 plates using several strategies to reduce the impact of technical artifacts. First, all longitudinal samples from the same patient were included on the same chip within the same plate so that longitudinal changes in methylation were not obscured by chip and plate effects. Second, row and column positioning of samples from the same patient were carefully assigned to available positions within a chip so that longitudinal changes in methylation were not confounded with row and column effects. Third, cases and controls for DCI were balanced within chips using a checkerboard pattern so that DCI was not confounded with row, column, chip, or plate effects (see Supplemental Figure 1 for the plate map). To gauge technical variation, we included four control samples of fixed methylation state (0%, 30%, 70%, and 100% methylated) and four technical replicates (i.e., repeated assays of the same DNA sample) per plate. Two of the control samples were placed in the same position across all plates and two were randomly placed. For the plate of blood samples, all four technical replicates were randomly positioned duplicates. In contrast, for the 11 plates of CSF samples, three of the four technical replicates were randomly chosen duplicate samples, and one was the same sample replicated across all 11 plates.

### 2.4 Sample Quality Functional Normalization

ENmix (Xu et al. 2016) was employed to assess the quality of samples in our methylation study, separately for blood and CSF samples. Samples having bisulphite control intensities less than 3 standard deviations below the mean of all samples, and/or for which more than 1% of probes were inadequately detected (i.e., detection p-values > 0.01 or with fewer than 3 beads) were categorized as low-quality samples. These, along with outliers in total intensity or beta value distribution were removed from our subsequent analyses (Xu et al. 2016). After the removal of low-quality and outlier samples, we performed background correction (Xu et al. 2016) to remove non-specific signals from the total signal, and performed dye bias correction (Xu et al. 2017). Sample quality differences by tissue type were tested using Fisher’s exact test on counts of samples passing or failing all sample QC filters.

We normalized the methylation data to bring Infinium Type I and Type II probes into alignment and to reduce noise and technical variation due to batch effects (i.e., plate, chip, row, and column effects). Specifically, we performed functional normalization, an extension of quantile normalization, which makes use of the control probes on the array to regress out unwanted variation in the methylation data (Fortin et al. 2014). Whether functional normalization improved agreement between technical replicates was tested by comparing the squared differences in median M-values between technical duplicates before and after normalization using a one-sided paired t-test.

### 2.5 CpG Site-Level Quality Control

After normalizing the data, we removed CpG sites from our analysis due to: (1) overlap of methylation probes with known polymorphic sites (which can cause biased methylation assessments), (2) probes located on the sex chromosomes (to rectify the artifacts arising due to unequal distribution of gender in the data) (Marabita et al. 2013), (3) cross-reactive probes that bind to alternate genomic sequences, (4) probes exhibiting multi-modal distributions indicative of poor quality or bias (Xu et al. 2016) and (5) probes that were inadequately detected (i.e., detection p-values > 0.01 or with fewer than 3 beads) in more than 1% of samples. Differences in the number of CpGs passing quality filters was tested using McNemar’s test.

### 2.6 Reference Based Cell Proportions for Blood

Blood has a mixture of cell types and DNA methylation-based references have been established for blood cells. Therefore, to estimate the proportions (cell counts) of each cell type, we employed Houseman’s reference based method (Houseman et al. 2012) using the functions available in the minfi package (Aryee et al. 2014) in our blood data. The method is based on using DNA methylation as a surrogate measure for cell type distributions and outputs the proportion of cell types: CD4+ T cells, CD8+ T cells, natural killer cells, monocytes, B -cells and granulocytes in each sample. The proportion of all cell types equals to one for each sample.

### 2.7 Cell-Type Heterogeneity Correction and Simulated EWAS Under the Null Hypothesis

Owing to the lack of reference methylation data for cell types found in CSF after an aSAH event we employed surrogate variable analysis (SVA) to perform reference-free adjustment for cell-type heterogeneity across the samples in blood and CSF data. SVA, as implemented in the sva R package (Leek et al. 2012), simultaneously models the effects of known sources of variation (covariates) and unknown sources of variation (i.e., surrogate variables), conditional on a phenotype of interest. Including the phenotype of interest in this modeling approach is necessary to prevent the surrogate variables from accounting for variation due to, for example, differences between cases and controls of disease, so as not to stymie subsequent analyses aimed at detecting CpG sites associated with case/control status. For examining the utility of surrogate variables in adjusting for cell-type heterogeneity in the absence of any particular phenotype-specific analyses, we generated a random trait by randomly permuting one of our observed traits, DCI, to serve as our outcome of interest. SVA was performed for this simulated trait along with age and gender as covariates in the context of an EWAS, whereby each CpG was individually tested for association with the simulated trait. Given the repeated measures in CSF, we grouped the CSF samples into five subsets centered on their target days (days 1, 4, 7, 10 and 13) and substituted samples +/- 1 day when a sample on the target day was unavailable. The goal of performing SVA cross-sectionally in CSF subsets is to retain the variation in methylation related to time. EWAS was also performed for the simulated trait without adjusting for surrogate variables and the distribution of p-values for SVA-adjusted and unadjusted EWAS scans under the null hypothesis were qualitatively compared to determine effect of SVA on genomic inflation. We measured inflation/deflation using the genomic inflation factor (**λ**), which is defined as the ratio of the empirically observed to expected median of the distribution of the test statistic.

### 2.8 Comparisons of Blood and CSF Methylation Profiles

We compared the methylation profiles of individuals with blood samples collected within the first day after hospitalization and CSF samples collected at day 1, 4, 7, 10 and 13. We used 65, 64, 65, 61 and 47 subjects to compare the methylation profiles of blood and CSF at day 1, day 4, day 7, day 10 and day 13 respectively to facilitate individual level comparison. For this comparison, we excluded CpG sites with a methylation beta value less than 10% or greater than 90% from all CpGs that passed QC, as methylation at these sites had little variation across samples and would not be informative for the analysis. The M-values at each qualifying CpG site were adjusted for age, sex and the surrogate variables using the aforementioned random trait to remove unwanted variation, and were then used to calculate correlation coefficients between the blood and CSF profile.

## 3 Results

### 3.1 Sample-level Quality Control

A total of 1,012 methylation profiles (including 44 technical replicates) were measured from CSF samples collected longitudinally from 279 aSAH patients. Additionally, 92 methylation profiles (including 4 technical replicates) were measured on blood samples in a subset of 88 of these patients; the majority of these blood samples (77) were sampled between zero and two days post-hospitalization (Table S1). QC analyses and filtering procedures were performed separately for CSF and blood samples. Based on low average bisulphite intensity and/or high proportion of poorly detected probes, we identified 89 (of 1012; 8.8%) poorly performing CSF samples (Figure 1). Additionally, we identified 3 (0.3%) more CSF outliers based on low total intensity. In contrast, no blood samples (0 of 92; 0%) failed these criteria. Figure 2 displays the beta-value distributions of all samples collected, based on which one blood and one additional CSF samples were identified as outliers. In total, poor sample performance was more common for CSF (93 of 1,012, 9.1%) than for blood (1 of 92, 1.1%), and these differences in quality of methylomic profiling by tissue type were statistically significant (Fisher’s exact test p = 0.003). Table S1 gives counts of all samples collected and samples retained after QC, for each collection time day.

**Figure 1:**
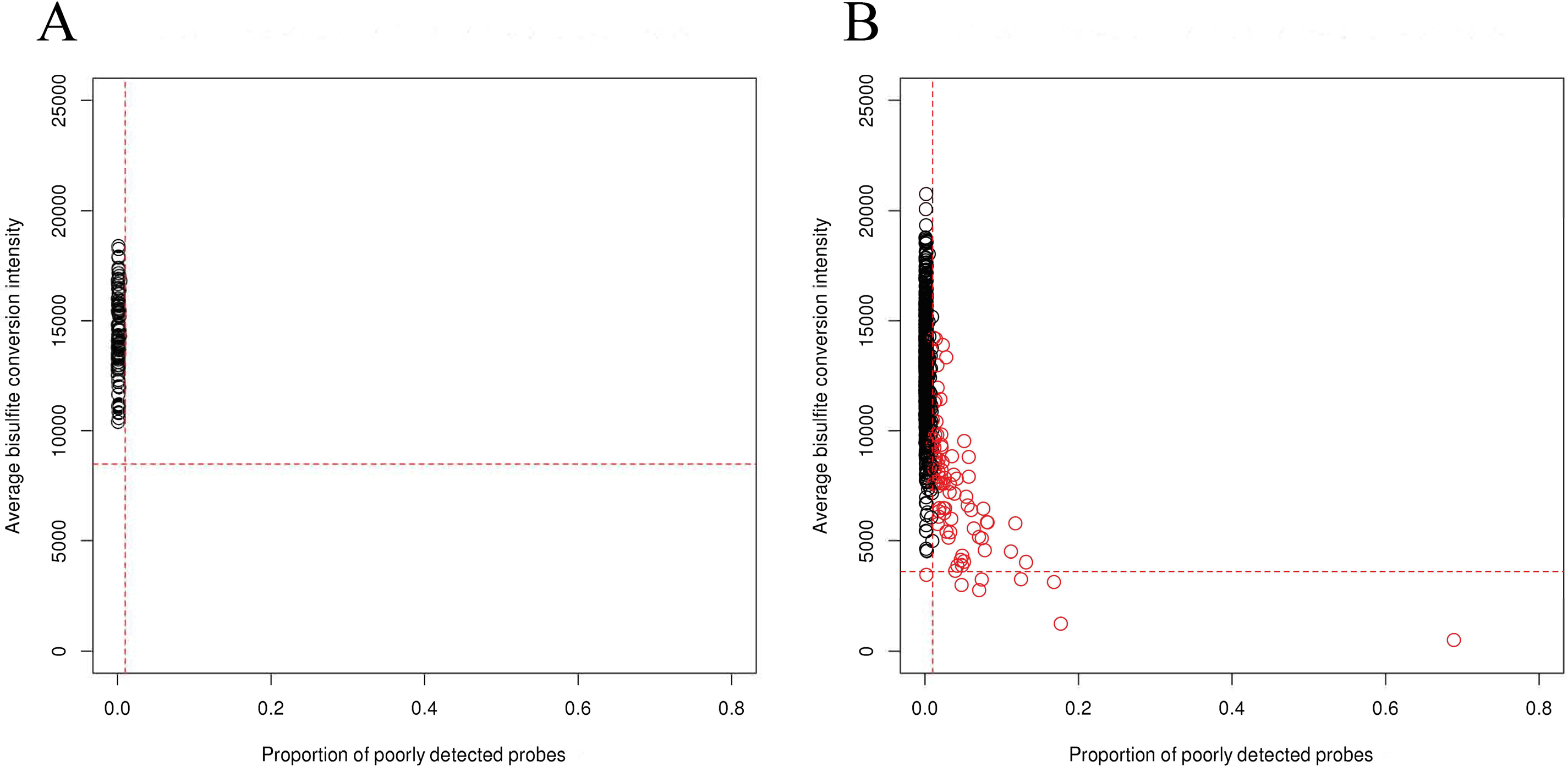
Identification of low-quality samples. (red) based on high proportion of poorly detected probes (x-axis) and/or low average bisulfite intensity (y-axis) from **(A)** 92 blood samples and **(B)** 1,012 CSF samples, both including technical replicates. The horizontal lines represent the threshold 3 SD below the mean across samples for bisulphite intensity, and the vertical lines represent the threshold of 1% of probes for which detection was poor (based on detection p-value and number of beads).

**Figure 2:**
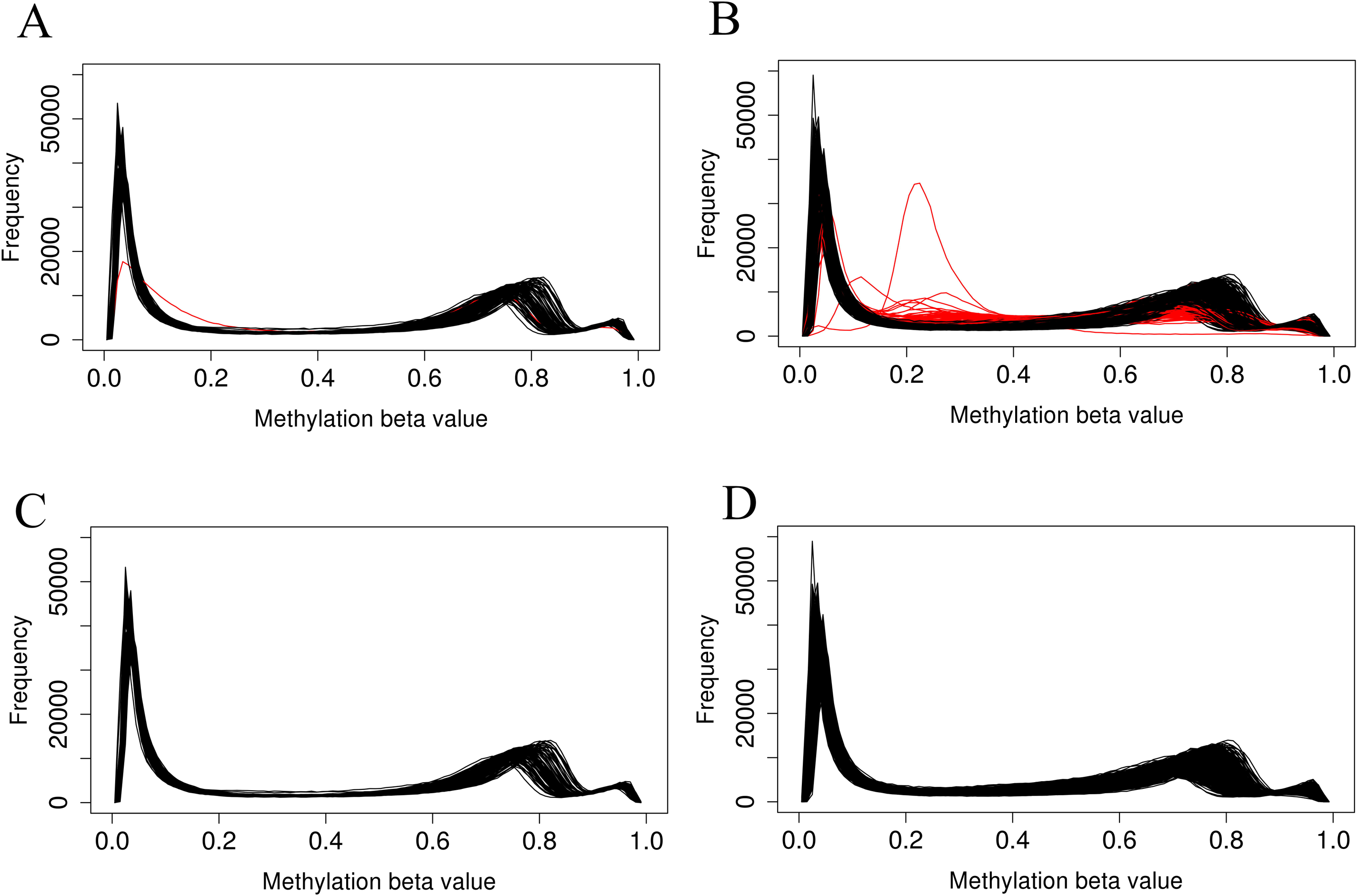
Distribution of beta-values across **(A)** all blood and **(B)** all CSF samples shows that a subset of poorly performing samples (red) deviate from the typical distribution. After removal of poor performing samples, distributions in **(C)** blood and **(D)** CSF are more consistent.

After removing low-quality samples, we performed functional normalization to reduce probe type (Infinium Type I vs. Type II) and batch (i.e., plate, chip, row, and column) effects. The reduction in chip, row, and column effects can be visualized in the distribution of M-values, before and after functional normalization, for samples profiled together on a plate (Figure 3). Row effects are apparent for some chips as increasing means across adjacent samples. For example, before normalization the third chip from the left in Figure 3A shows strong row effects indicated by means forming an upwardly sloped trend across the first to fifth samples (which correspond to ascending rows in the first column), followed by another upwardly sloped trend across the sixth to eleventh samples (which correspond to ascending rows in the second column). Functional normalization increased concordance in median methylation between 34 technical replicate CSF samples (p = 0.015) (Figure 4). For the 4 technical replicate blood samples, the same trend of increased concordance after functional normalization was observed; however, this trend was not statistically significant (p = 0.153).

**Figure 3:**
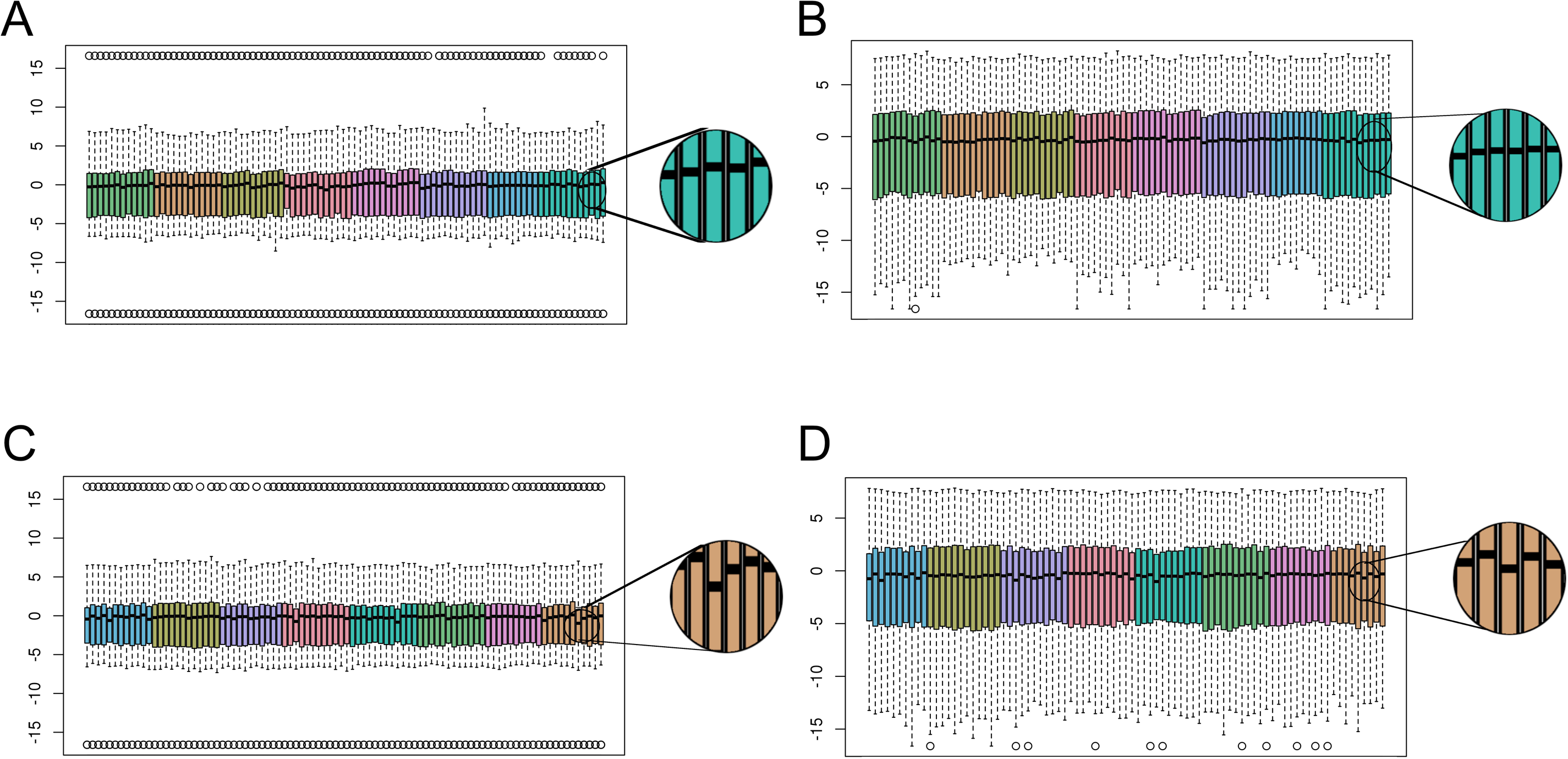
Functional normalization reduces batch effects. Boxplots show the distribution of M-values per sample across the plate of **(A)** blood samples before, and **(B)** after, functional normalization. For each sample, the median M-value is indicated by the black horizontal line and the interquartile range (25^th^ to 75% percentile) is indicated by the colored box. The whiskers (dashed lines) extend to the most extreme data point within 1.5 times the interquartile range beyond the box, and outlier points beyond this limit are shown individually as circles. Samples are colored coded by chip, and samples are ordered within each chip as follows: first column ascending by row number followed by second column ascending by row number. Before normalization, chip effects are apparent as differences in median and interquartile range between color groups. **(C)** CSF samples before, and **(D)** after, functional normalization on an example plate. Comparing the blow-ups to the right of each plot show variation in median M-values across samples is reduced after functional normalization.

**Figure 4:**
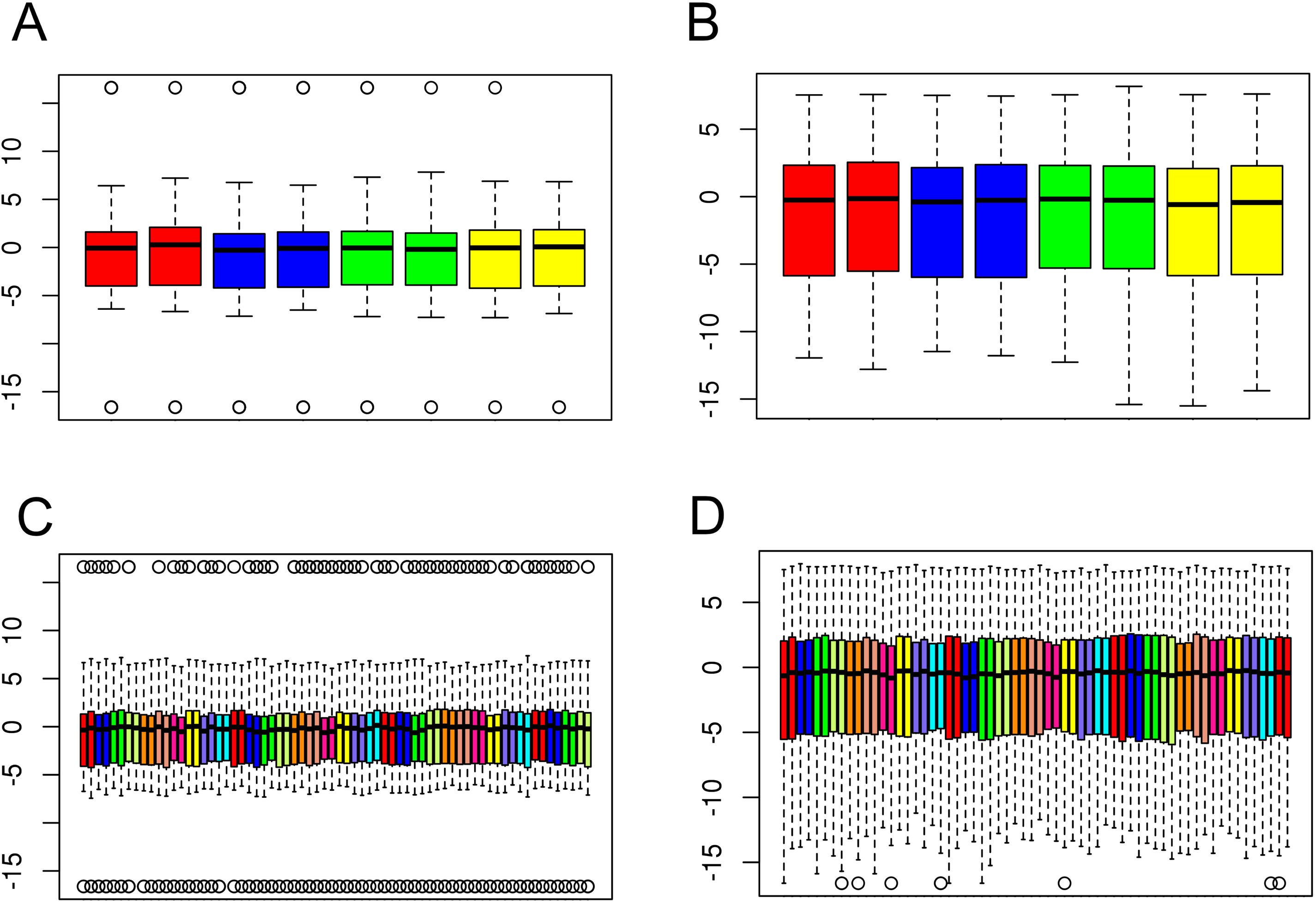
Functional normalization increases concordance of technical replicates. Boxplots showing distribution of M-values for duplicate **(A, B)** blood and **(C, D)** CSF samples **(A, C)** before, and **(B, D)** after, functional normalization. Pairs of duplicates are adjacent to each other and differentiated by color. For each sample, the median M-value is indicated by the black horizontal line and the interquartile range (25^th^ to 75^th^ percentile) is indicated by the colored box. The whiskers (dashed lines) extend to the most extreme data point within 1.5 times the interquartile range beyond the box, and outlier points beyond this limit are shown individually as circles.

### 3.2 CpG probe-level Quality Control

Individual probes were filtered out of analyses for reasons pertaining to probe design such as overlap with common single nucleotide polymorphisms (SNPs) and cross-reactivity with off-target genomic positions. Additionally, CpG probes on the sex-chromosomes were excluded. Based on QC analyses, CpG probes with multimodal beta-value distributions, low detection quality across samples, and high technical variation across replicate samples were also filtered out of analyses. CpG probe-level filtering criteria are summarized in Table 2. For each QC filtering step, and overall, fewer CpGs were filtered out in blood than in CSF (p < 2.2 × 10^−16^ for all), indicating that CSF samples may yield somewhat lower-quality methylation data, as is also evident in Figure 1.

**Table 2:**
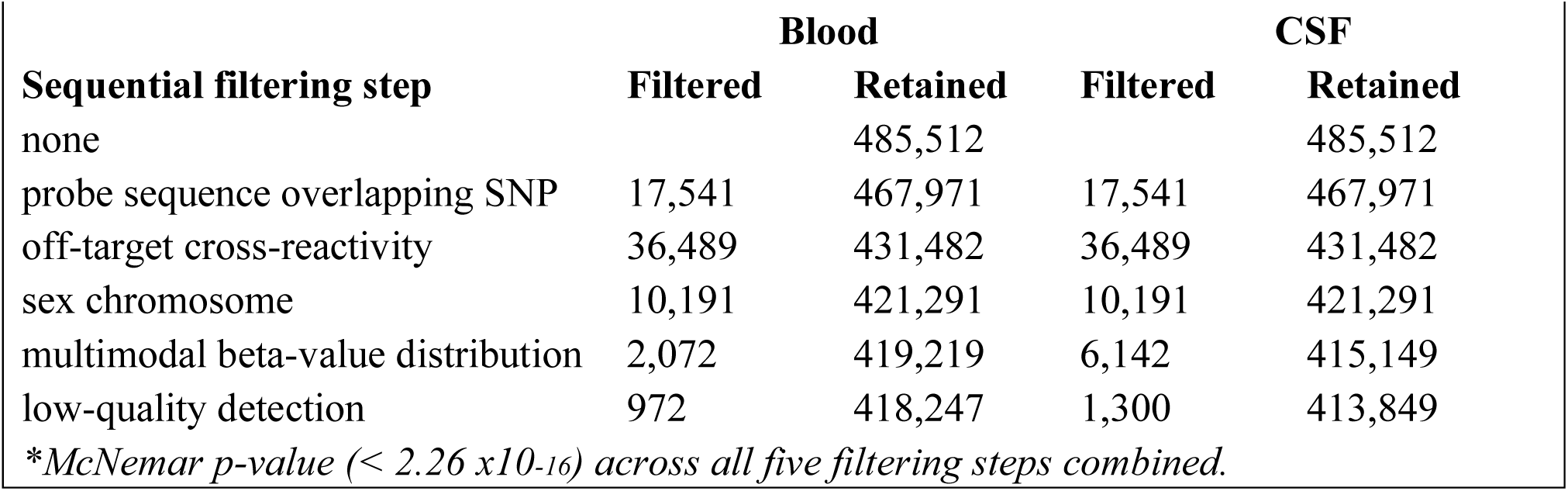
CpG probe-level filters.

| Sequential filtering step | Blood |  | CSF |  |
| --- | --- | --- | --- | --- |
|  | Filtered | Retained | Filtered | Retained |
| none |  | 485,512 |  | 485,512 |
| probe sequence overlapping SNP | 17,541 | 467,971 | 17,541 | 467,971 |
| off-target cross-reactivity | 36,489 | 431,482 | 36,489 | 431,482 |
| sex chromosome | 10,191 | 421,291 | 10,191 | 421,291 |
| multimodal beta-value distribution | 2,072 | 419,219 | 6,142 | 415,149 |
| low-quality detection | 972 | 418,247 | 1,300 | 413,849 |

*\*McNemar p-value ( $< 2.26 \times 10^{-16}$ ) across all five filtering steps combined.*

### 3.3 B-cell Leukemia Outlier

Estimated blood cell type proportions using the reference-based method followed expectations for all blood samples with one exception, which showed high B-cell composition in analysis. Further clinical investigation confirmed the presence of chronic lymphocytic leukemia (CLL) in the individual, which is known to cause increased proliferation of B cells in blood, bone marrow and other lymphoid tissues (Zhang and Kipps 2014; Ciccone et al. 2014; Hallek 2015; Greenberg and Probst 2013; Ghia and Hallek 2014). Samples from this participant were excluded from further analyses.

### 3.4 Adjustment for Cell Type Heterogeneity

Because methylomic profiles differ widely by cell type, modeling cell type heterogeneity across samples is crucial for valid cross-sample analyses of methylation data. However, external cell type-specific reference data was not available for post-aSAH CSF for use in reference-based adjustment. Therefore, we performed reference-free adjustment using SVA to remove unknown sources of variation including cell type heterogeneity. We further excluded technical replicates from all samples that passed QC, leaving 70 blood samples and 154, 246, 217, 152, and 95 CSF samples for days 1, 4, 7, 10 and 13 respectively. Ten surrogate variables (SVs) were generated for the set of blood samples, and 13 SVs were generated for day 1 CSF samples. Fifteen, 15, 14 and 10 surrogate variables were generated for CSF samples for day 4, 7, 10 and 13 respectively. To determine the benefit of SV-adjustment, we interrogated its effect on CpG site association tests under the null model of no association by simulating a dummy binary phenotype similar to the distribution of DCI and performing EWAS, with and without including SVs as covariates. The behavior of the test statistic better followed the null distribution after SV-adjustment, as shown in quantile-quantile plots (Figure 5). Specifically, genomic inflation factor (**λ**) improved from 1.11 to 0.98 in the set of blood samples, and improved from 0.73 to 0.99 in the set of CSF samples within two days after hemorrhage (Figure 5) and likewise in other CSF subsets (Supplemental Figures 2 and 3). Genomic deflation may be caused by sources of variation including cell type heterogeneity that cause correlation across CpG sites within a sample, equating to a reduction in the effective number of independent tests. These results show that in the absence of reference data, SVA aids in controlling the adverse impact of cell-type heterogeneity and other sources of unwanted variation on tests of epigenetic association.

**Figure 5:**
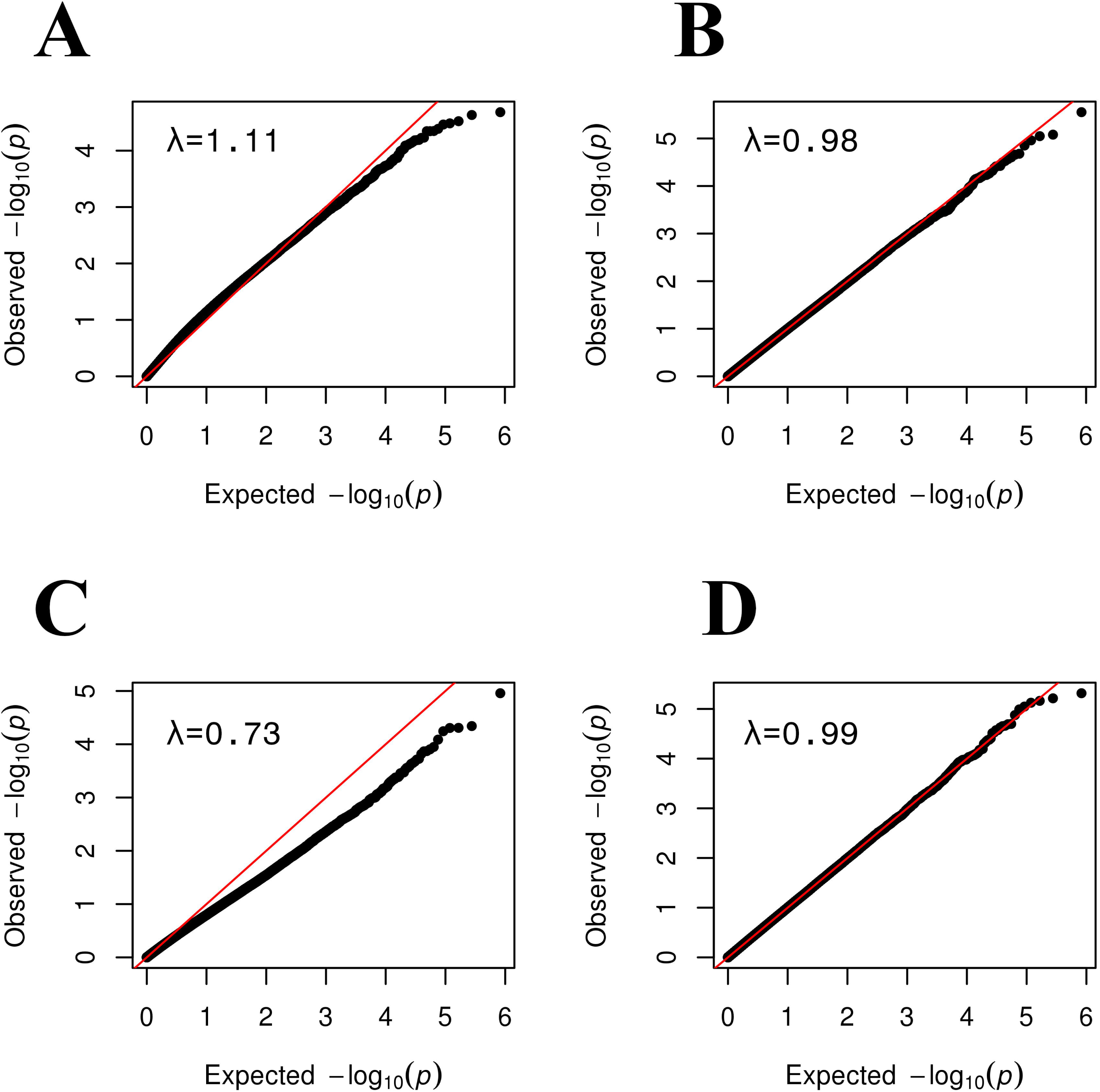
Quantile-quantile plots showing the benefit of SVA for tests of epigenetic association. The distribution of observed p-values obtained for a random simulated phenotype (y-axis) are plotted against the expected distribution of p-values under the null model of no association in (A, B) blood samples, and (C, D) CSF samples at day 1. The genomic inflation factor, λ, is at top left on each plot. (A) Simulated EWAS without SV-adjustment showed inflation with a λ =1.11 (B) After SV-adjustment the EWAS closely follows the null distribution as indicated by points closely following the diagonal. **(C)** Simulated EWAS exhibits genomic deflation with a λ = 0.73 (D) After SV-adjustment, the EWAS closely follows the null distribution (i.e., points closely following the diagonal).

### 3.5 Correlation was low when comparing DNA methylation of post-aSAH blood and CSF

Following our long-term goal of understanding the methylomic changes occurring across tissues after aSAH, we explored the suitability of peripheral blood collected within the first day of hospitalization as a surrogate for the normally less accessible longitudinally collected CSF based on the correlation of adjusted M-values between the two tissue types obtained from aSAH patients. Specifically, we compared the methylation profile of blood collected within 48h of hospitalization versus CSF samples collected at day 1, 4, 7, 10 and 13 post rupture respectively. Table 3 summarizes the numbers of CpGs used and the correlation coefficients for each day. In general, the mean correlation (0.23 - 0.26) was too low to use blood as a surrogate for post-aSAH CSF in a global manner.

**Table 3:**
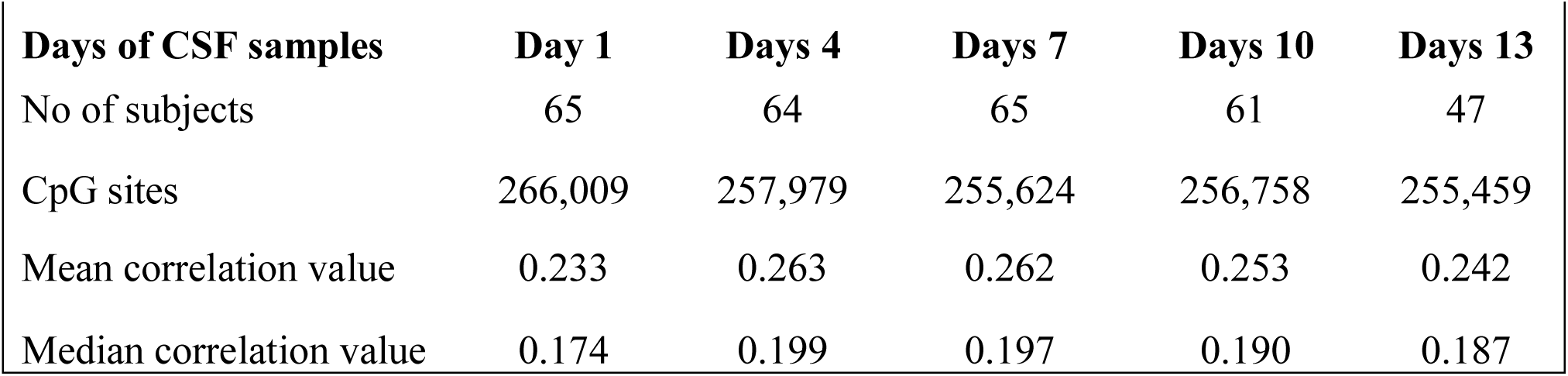
Correlation analysis of blood (within first day of hospitalization) and CSF at different times

| Days of CSF samples | Day 1 | Days 4 | Days 7 | Days 10 | Days 13 |
| --- | --- | --- | --- | --- | --- |
| No of subjects | 65 | 64 | 65 | 61 | 47 |
| CpG sites | 266,009 | 257,979 | 255,624 | 256,758 | 255,459 |
| Mean correlation value | 0.233 | 0.263 | 0.262 | 0.253 | 0.242 |
| Median correlation value | 0.174 | 0.199 | 0.197 | 0.190 | 0.187 |

Differences were observed in the magnitude of the correlation by genomic position (CpG island, shore, shelf, and open sea; p < 0.001 for all), with islands and shores showing greater positive correlation than shelves and seas (Figure 6, Supplemental Figures 4-7). Similarly, the magnitude of the correlation differed by the orientation of CpG with respect to the nearest gene ([3’ UTR, TSS, Exon, Body, 5’UTR], p < 0.01), with CpG sites near the transcription start site or first exon showing greater inter-tissue correlation than CpG sites in the upstream, downstream or in the body of genes. The CpGs sites upstream or in the body of genes, in turn, showed greater correlation than CpG sites downstream of the gene.

**Figure 6:**
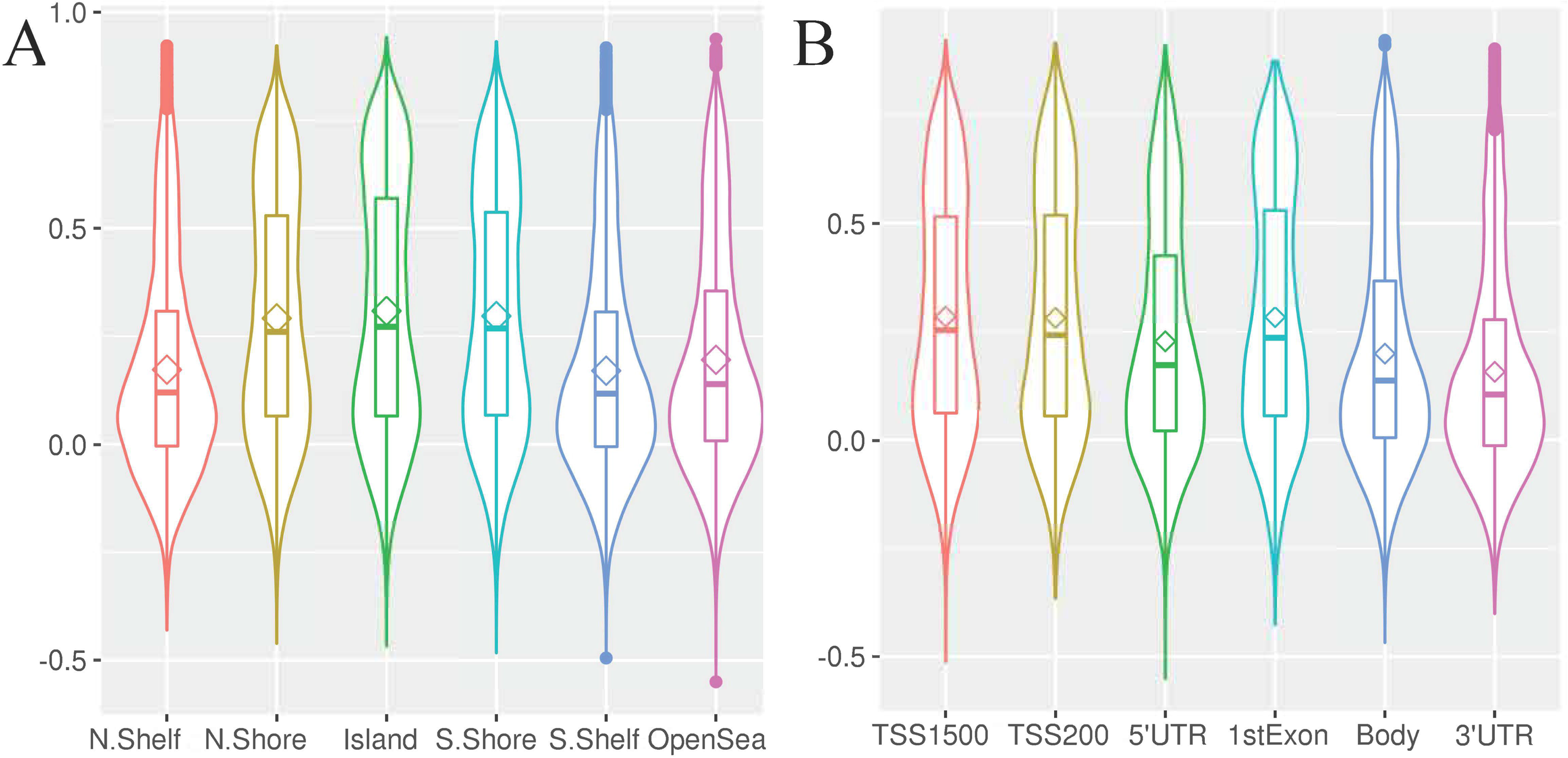
Correlation between blood and CSF at day 1 for CpG sites across **(A)** genomic regions, and **(B)** relative to genes. Bean plots depict the median correlation coefficient (horizontal line), mean (diamond), interquartile range (i.e., 25^th^ to 75^th^ percentile, box), and density (width of the bean). (TSS – Transcription start site; UTR – Untranslated region).

## 4 Discussion

Our protocol demonstrated the value of several QC procedures in obtaining clean and useful methylation data for subsequent scientific analyses. In particular, in addition to quality filters at the sample and CpG probe level, we showed that functional normalization was helpful in reducing batch effects for both blood and CSF. Likewise, SVA was useful for adjusting for unknown sources of variation, including cell type heterogeneity, as evidenced by improved genomic inflation factor for a simulated EWAS scan under the null hypothesis. This observation is particularly important for studies of tissue types, such as CSF, that are underrepresented in the methylomics literature, and for which external cell type reference data are not yet available. We also provided evidence that, overall, CSF samples yielded lower-quality methylomic data than did blood samples. This observation may reflect the low cell content (de Graaf, Smitt, et al. 2011; de Graaf, de Jongste, et al. 2011; Svenningsson et al. 1995) in CSF compared to blood. Altogether, these lessons can inform the design of future analyses seeking to investigate the methylomic profiles in post-aSAH CSF samples. The efficiency of a reference-based method in capturing the outlier with high proportion of B-cells is promising.

We also explored the question of whether methylomic profiles from blood samples could serve as surrogates for less accessible CSF. Though significant positive correlations were observed, especially for regulatory regions such as CpG islands and locations near transcriptional start sites of genes, globally, the correlations in methylation values between blood and CSF were too low for blood to serve as a useful surrogate for most scientific or clinical purposes. However, to understand the methylomic changes that occur post-aSAH, we believe that CSF would be a most relevant source, representing the central nervous system (CNS) environment and its proximity to the hemorrhagic location.

This study benefited from several strengths including the thoughtful plate design aimed at reducing confounding of experimental effects with technical artifacts, thorough and rigorous application of data QC procedures, pairing of blood and CSF samples from the same patients, and assessment of methylomic profiles in a novel tissue type (post-aSAH CSF) that captures the CNS environment post-aSAH. The study is also novel as this is the first to investigate methylation patterns in DNA extracted from CSF over a longitudinal period after aSAH. Despite these strengths, limitations of the current study include limited statistical power to resolve the intra subject differences among the samples that may ultimately pose challenges in using this dataset for future EWAS studies. Additionally, the cell composition of CSF may vary over time after hemorrhage, which would also affect the methylation levels. Thus, longitudinal analyses of post-aSAH samples are challenging as cell-type heterogeneity may be confounded with days post injury. Overcoming these challenges will be necessary to accomplish goals such as identifying genes whose changes in methylation after injury are predictive of recovery outcomes.

In conclusion, this study is one of the first attempts to investigate DNA methylation at the genome scale in a sample of aSAH patients, as well as one of the first to measure methylation in CSF. Our analysis protocol showed that methylomic profiles can be obtained from CSF for use in EWAS analysis and that QC steps can improve the analysis by eliminating low-quality data points and reducing biases and experimental artifacts. Likewise, we show that blood, while readily accessible, is not a sufficient surrogate for the methylomic status of CSF. Ultimately, efforts to understand methylation profiles in aSAH patients, and changes that occur post-injury, may lead to the discovery of biomarkers of clinical utility in predicting patient recovery.

## Author contributions

YC is the principal investigator of the project. YC, DW and JS conceived and designed the study. AA, DL, TK, JS, DW performed the experiments and statistical analysis. AIA, JRS and DEW contributed to the initial writing of the manuscript. EAC contributed to the clinical investigation of the research. All authors reviewed, edited, and approved the final manuscript.

## Acknowledgements

Foremost, we thank the participants of this study for making this work possible. Funding for this study was provided by the National Institute of Health (R01NR013610). We thank UPMC for access of the clinical samples.

## Public Data Access

The data can be accessed through dbGAP: phs001990.v1.p1.

