## Supplemental for "Methylation Data Processing Protocol & Comparison of Blood and Cerebral Spinal Fluid Following Aneurysmal Subarachnoid Hemorrhage"

**1 Supplementary Figures**

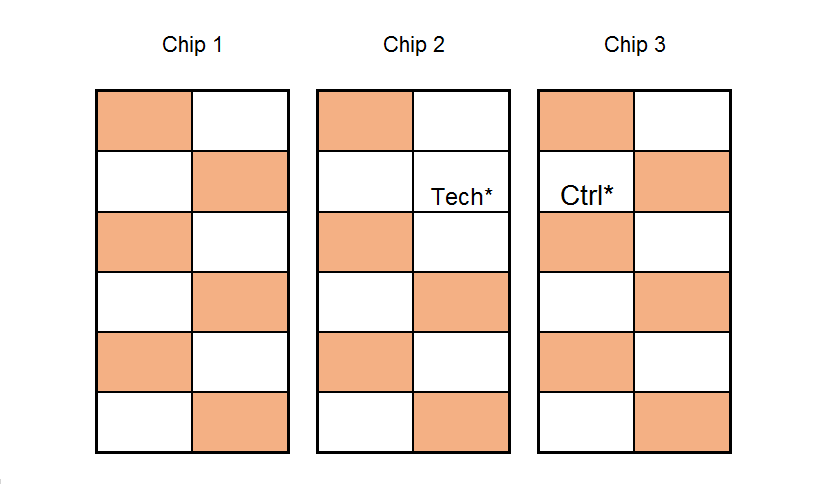

**Supplementary Figure 1.** A representative plate map. Cases (colored boxes) and controls were balanced within chips using a checkerboard pattern to reduce row, column, chip, or plate effects.

[Tech* denotes technical duplicates; Ctrl* denotes methylation controls].

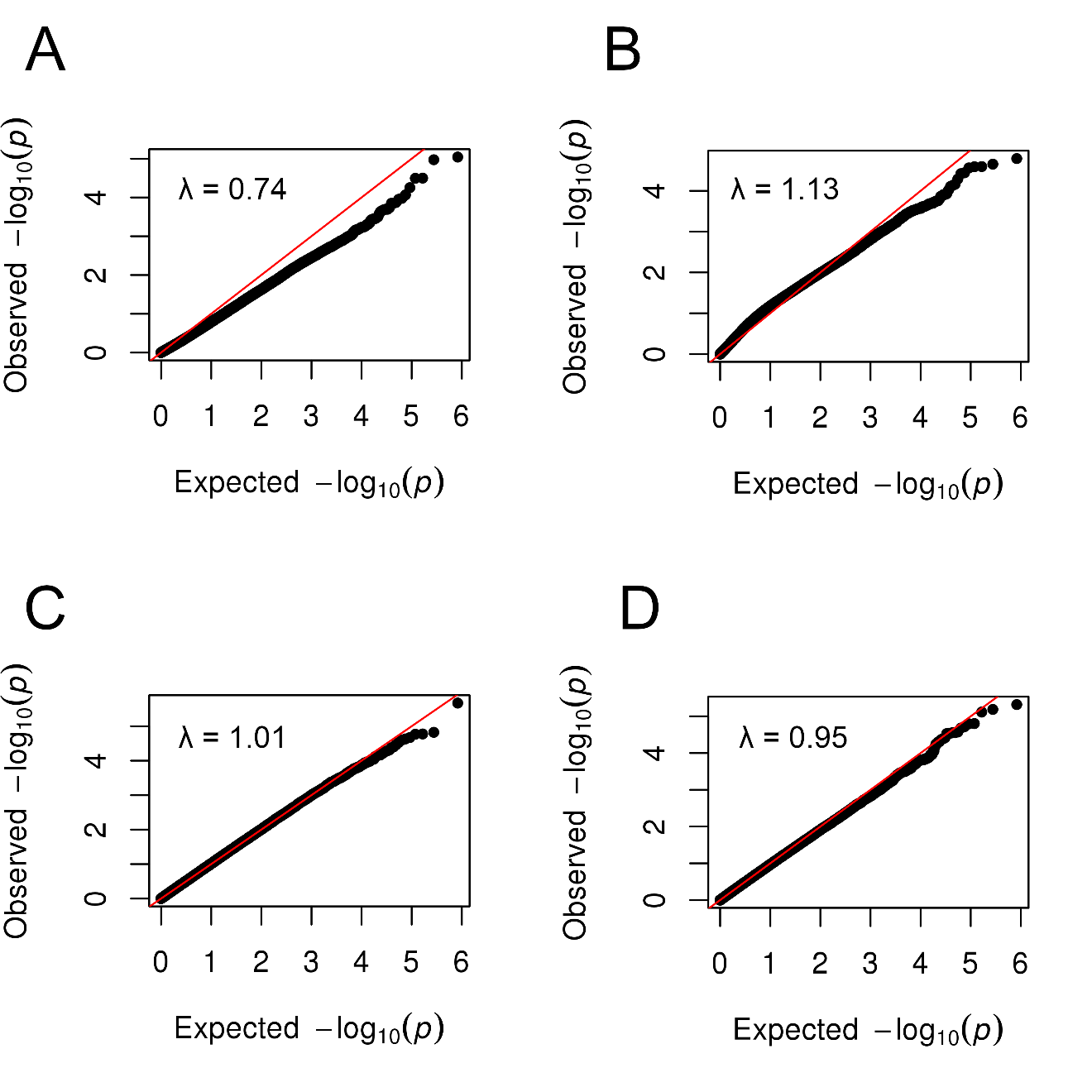

**Supplementary Figure 2.** The distribution of observed p-values obtained for a random simulated phenotype (y-axis) are plotted against the expected distribution of p-values under the null model of no association in CSF samples at day 4 (left) and day 7 (right) respectively. The upper panel shows the simulated EWAS without SV-adjustment (A, B) and the lower panel illustrates after SV-adjustment (C, D). The genomic inflation factor, **λ**, is indicated on each plot.

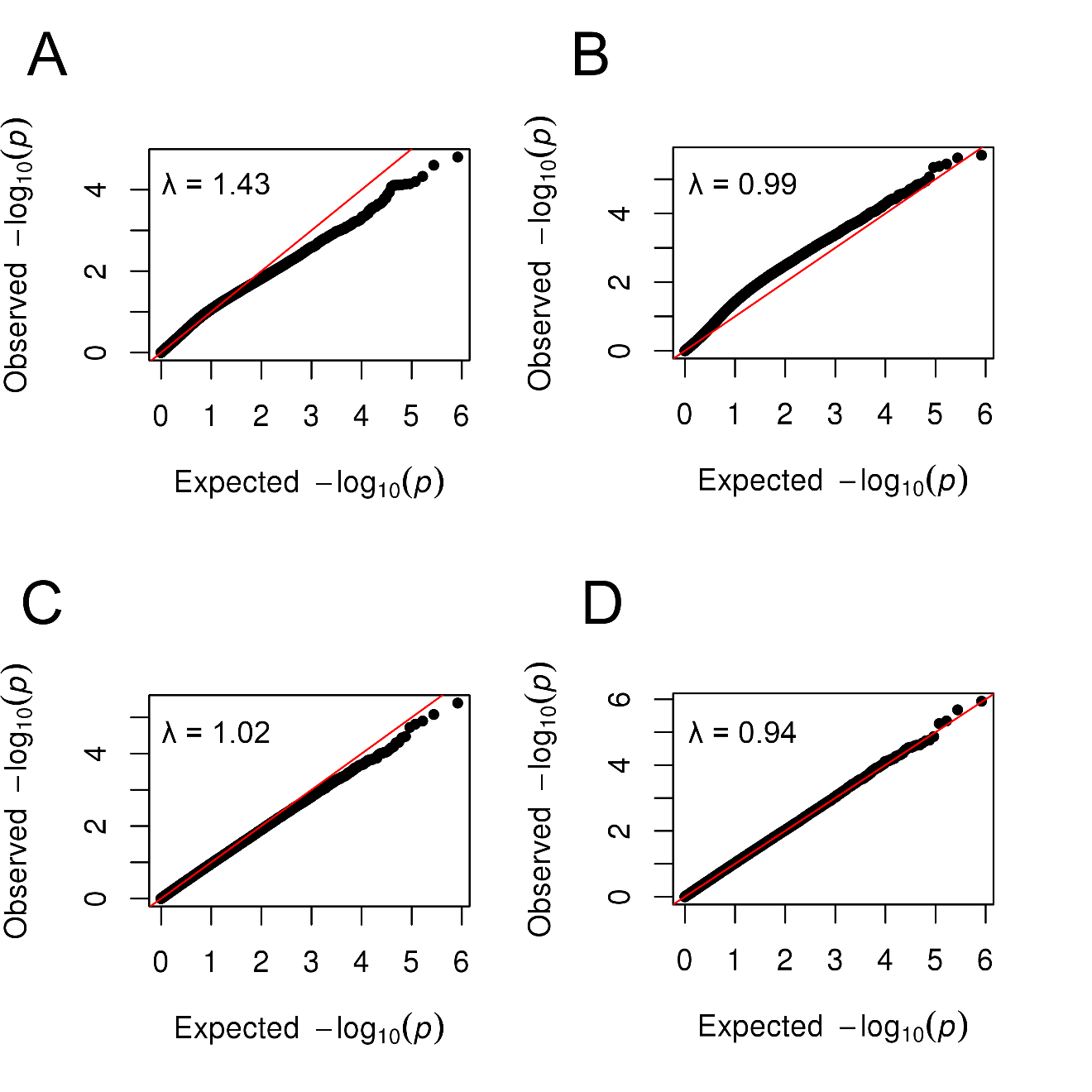

**Supplementary Figure 3.** The distribution of observed p-values obtained for a random simulated phenotype (y-axis) are plotted against the expected distribution of p-values under the null model of no association in CSF samples at day 10 (left) and day 13 (right) respectively. The upper panel shows the simulated EWAS without SV-adjustment (A, B) and the lower panel illustrates after SV-adjustment (C, D). The genomic inflation factor, **λ**, is indicated on each plot.

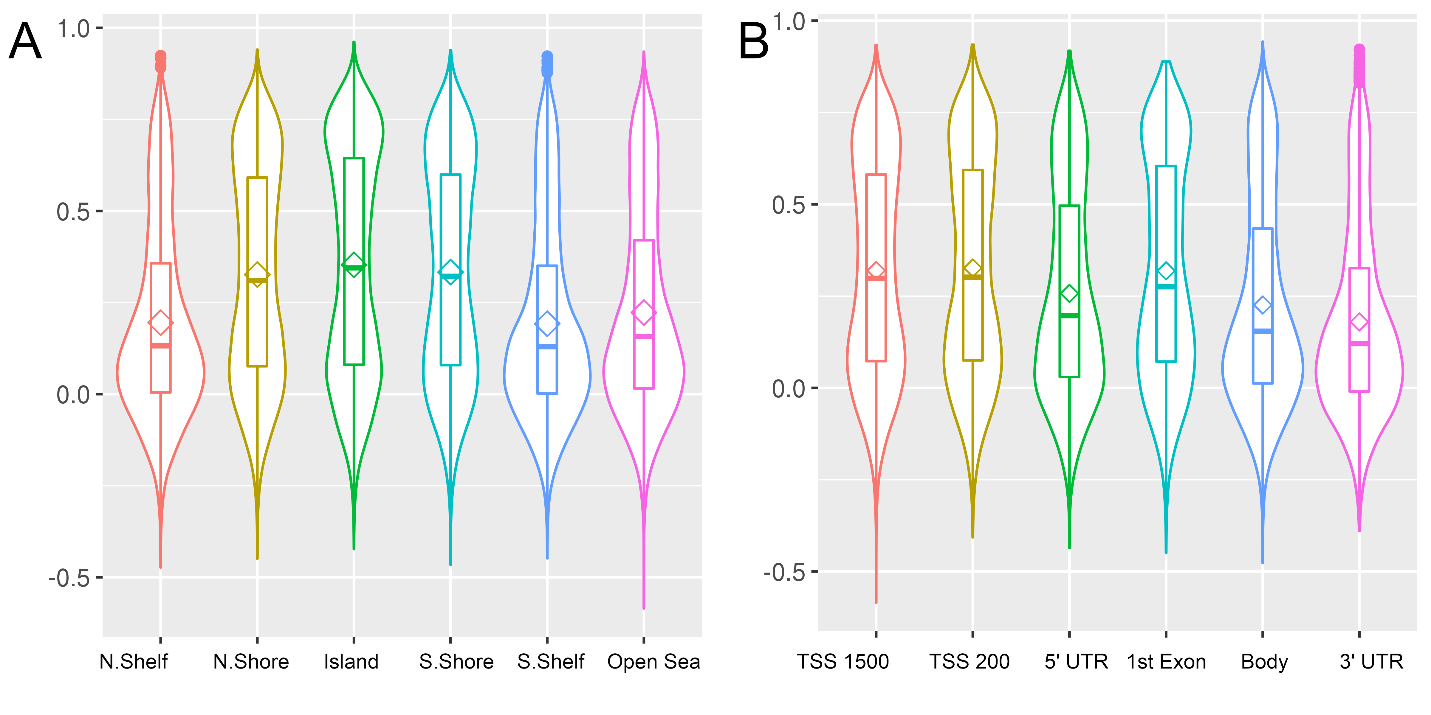

**Supplementary Figure 4.** Correlation between blood and CSF at day 4 for CpG sites across genomic regions (A), and relative to genes (B). Bean plots depict the median correlation coefficient (horizontal line), mean (diamond), interquartile range (i.e., 25^th^ to 75^th^ percentile, box), and density (width of the bean). TSS – Transcription start site; UTR – Untranslated region.

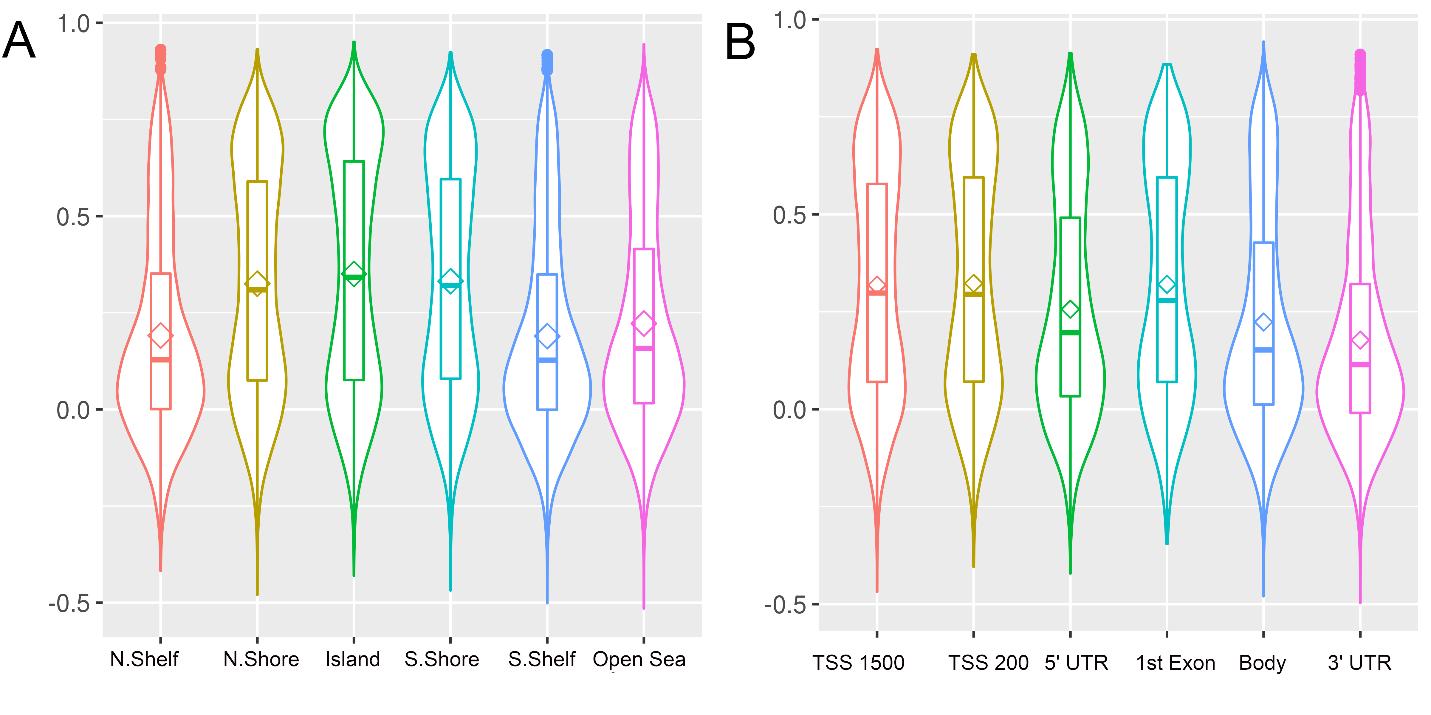

**Supplementary Figure 5.** Correlation between blood and CSF at day 7 for CpG sites across genomic regions (A), and relative to genes (B).

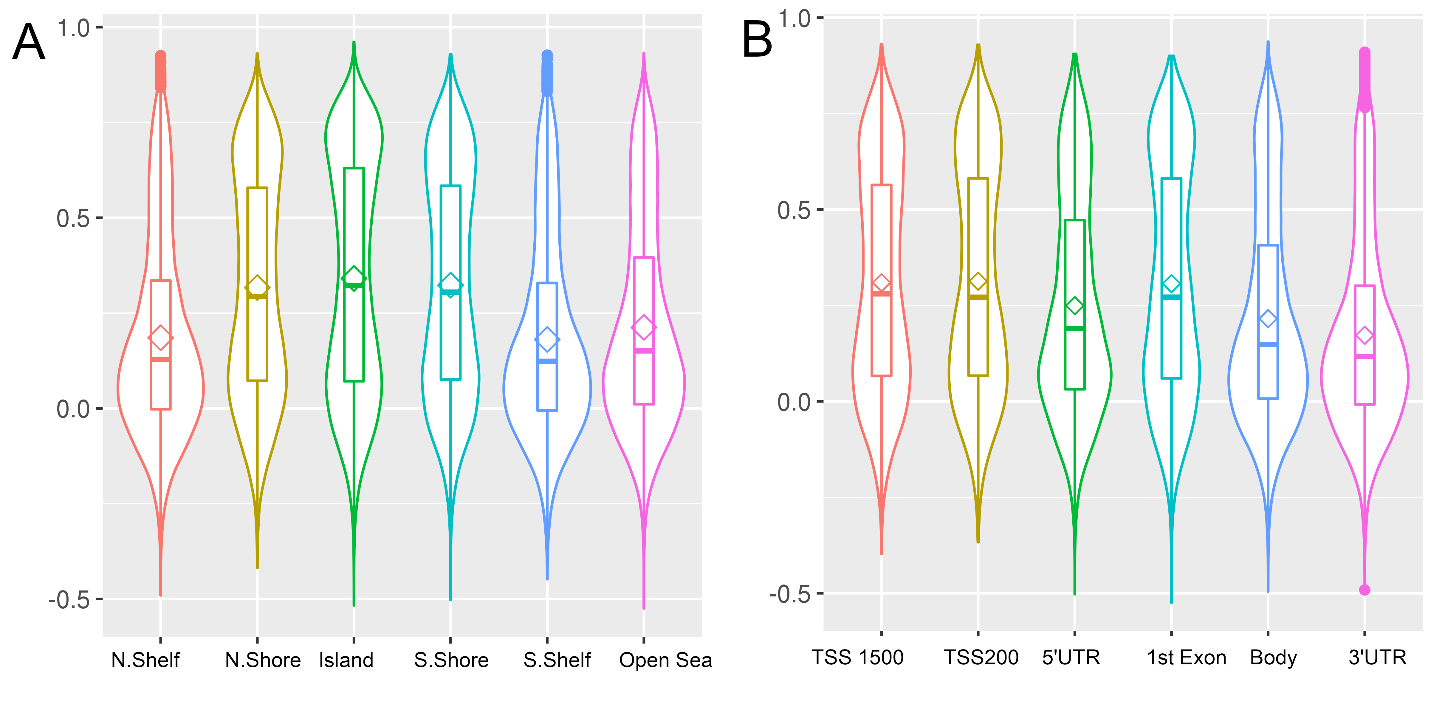

**Supplementary Figure 6.** Correlation between blood and CSF at day 10 for CpG sites across genomic regions (A), and relative to genes (B).

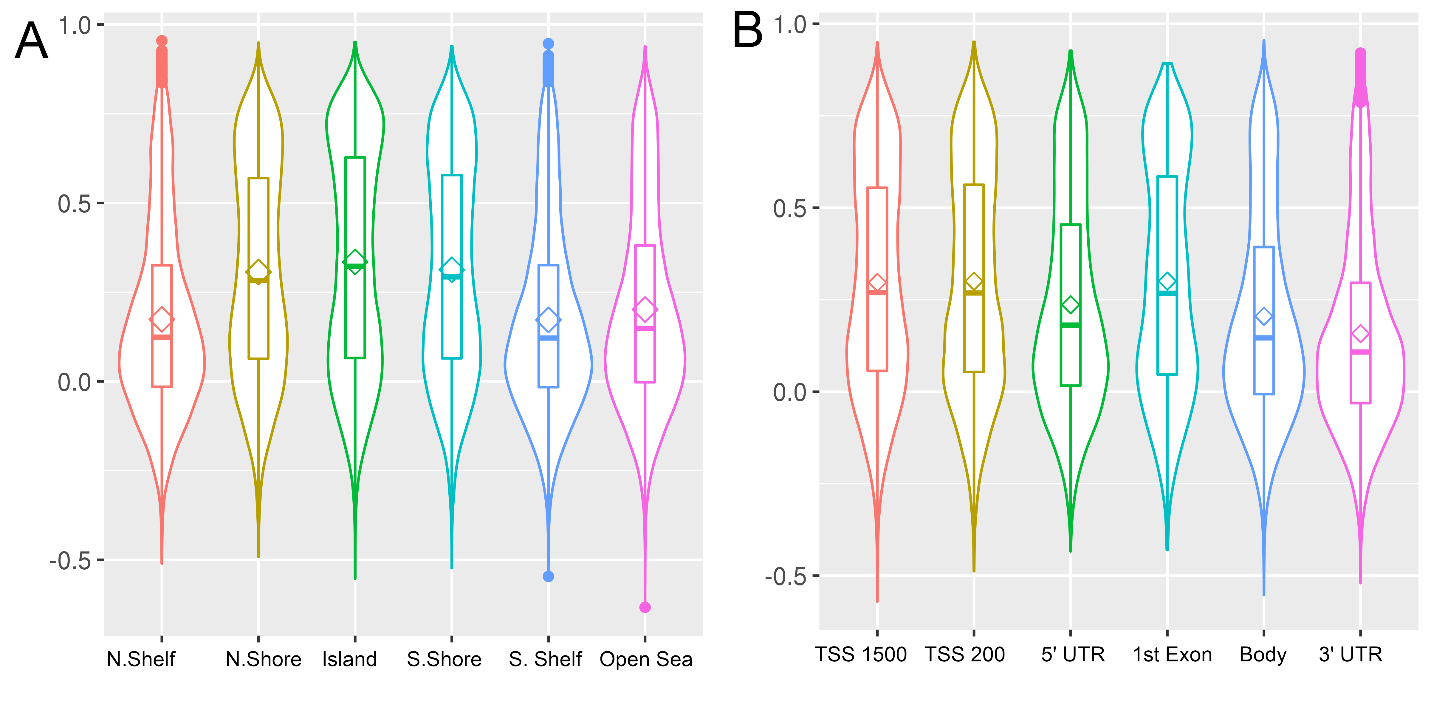

**Supplementary Figure 7.** Correlation between blood and CSF at day 13 for CpG sites across genomic regions (A), and relative to genes (B).

**2 Supplementary Table**

**Table S1**. Number of total samples collected and samples that passed quality control, at each target day and ±1 day

|  | **Total sample collected** | | | |  | **Sample passed QC** | | | |
| --- | --- | --- | --- | --- | --- | --- | --- | --- | --- |
| **CSF** |  |  |  |  |  |  |  |  |  |
| Day group | -1 | Target day ^a^ | +1 | Total |  | -1 | Target day | +1 | Total |
| 0-2 | 3 | 83 | 99 | 185 |  | 2 | 76 | 90 | 168 |
| 3-5 | 56 | 114 | 121 | 291 |  | 54 | 102 | 110 | 266 |
| 6-8 | 61 | 96 | 94 | 251 |  | 55 | 86 | 88 | 229 |
| 9-11 | 41 | 79 | 56 | 176 |  | 34 | 72 | 51 | 157 |
| 12-14 | 36 | 52 | 20 | 108 |  | 35 | 47 | 17 | 99 |
| 15 | 0 | 1 | 0 | 1 |  | 0 | 0 | 0 | 0 |
| **Total** |  |  |  | 1012 |  |  |  |  | 919 |
| **Blood** |  |  |  |  |  |  |  |  |  |
| Day group | -1 | Target day | +1 | Total |  | -1 | Target day | +1 | Total |
| 0-2 | 2 | 34 | 41 | 77 |  | 2 | 34 | 40 | 76 |
| 3-5 | 6 | 7 | 0 | 13 |  | 6 | 7 | 0 | 13 |
| 6-8 | 0 | 0 | 2 | 2 |  | 0 | 0 | 2 | 2 |
| **Total** |  |  |  | 92 |  |  |  |  | 91 |

^a^ Target days are 1,4,7,10 and 13
